# Human foregut morphogenesis exhibits murine-like molecular patterning but avian-like epithelial architecture

**DOI:** 10.64898/2026.09.17.752534

**Authors:** Rui Yan, Albert Z. Ma, Clifford J. Tabin

## Abstract

Animal models are indispensable for understanding human development, yet evolutionary relatedness does not ensure developmental similarity at all biological scales. In tracheal-esophageal separation (TES), which splits the embryonic foregut into respiratory and digestive tubes in all tetrapods, genetic mouse models rarely reproduce the predominant human malformation, esophageal atresia with tracheoesophageal fistula (EA/TEF). By comparing human, mouse, and chick foreguts, we uncover a mosaic pattern of developmental conservation. The human foregut resembles the mouse in its molecular patterning, but more closely resembles the chick in its densely pseudostratified epithelial architecture and the greater number of cells comprising the epithelial septum. Live imaging shows that epithelial pseudostratification, which depends on actomyosin contractility, is associated with slower septum resolution. Cross-species single-cell transcriptomics analysis reveals lower expression of a myosin regulatory light chain and N-cadherin, and higher expression of a myosin phosphatase subunit in the mouse foregut epithelium than in the human and chick, suggesting candidate molecular mechanisms of these morphological differences. Induction of ectopic BMP signaling through in vivo electroporation in chick embryos recapitulated an EA/TEF-like malformation. These findings suggest that human foregut morphogenesis combines murine-like patterning and avian-like epithelial organization. The chick embryo thus represents an important complementary model for investigating epithelial mechanisms underlying human foregut malformations.

## Introduction

The embryonic foregut gives rise to the upper digestive tract and the respiratory system in tetrapods through tracheal-esophageal separation (TES), a morphogenetic process that splits the foregut through epithelial tube constriction and septation (1, 2). Defects in foregut morphogenesis cause congenital malformations of the esophagus and trachea, affecting approximately 1 in 2,500 live births (3). Among these, esophageal atresia and tracheoesophageal fistula (EA/TEF), characterized by an occluded proximal esophagus pouch and an abnormal connection between the distal esophagus and the airway, is the most common (∼1 in 4,000) (4). The morphogenetic mechanism of EA/TEF remains poorly understood due to the lack of a dynamic view of TES, although early descriptive studies have revealed human foregut morphology in fixed embryonic tissues across developmental stages (5, 6).

Model organisms have proved useful in understanding the morphogenetic mechanisms of human organogenesis, but we do not know how faithfully commonly used models reproduce human TES and related congenital malformations. Although the core signaling pathways in foregut patterning are conserved across tetrapods (7), morphogenesis can differ significantly depending on species-specific tissue characteristics. Because mice are more closely related to humans than non-mammals, and the availability of exceptional genetic tools, the mouse model is most widely used. Paradoxically, however, very few genetic mouse models recapitulate EA/TEF. Rather, most mouse models with defective TES instead undergo complete separation failure, exhibiting laryngotracheoesophageal cleft (LTEC), which is much rarer in humans (1 in 40,000) (3, 4). The frog embryo is another established model system for studying TES, which has successfully recapitulated septation defects with human-related genetic perturbations (8, 9). However, the gross morphology of the frog foregut differs considerably from that of mammals, with a relatively short and narrow trachea. Recently, we established the chick embryo as a compelling model for TES, which allows for robust ex vivo culture, live imaging, and various mechanical and pharmacological perturbations of this dynamic process (10). Notably, chick and mouse foreguts show significant differences in molecular patterning and epithelial architecture (10). It remains unclear which features of foregut morphogenesis are evolutionarily conserved between the human and different experimental models, and whether these differences influence how genetic perturbations manifest as distinct congenital malformations.

Here, we compare foregut patterning and epithelial organization in fixed foregut sections from human, mouse, and chick embryos during TES. We find that although humans and mice share most molecular patterns, the human foregut epithelium is surprisingly similar to the chick in its architecture, both exhibiting a more pseudostratified organization than the relatively simple mouse foregut epithelium. Epithelial architecture, controlled by actomyosin forces, is associated with the dynamics of epithelial septum resolution. Integrating single-cell RNA-sequencing (scRNA-seq) datasets of human, mouse, and chick foreguts, we find species-specific gene expression profiles that may underlie differential epithelial organization. These findings suggest that the chick embryo may be useful for modeling tracheal-esophageal malformations arising from epithelial abnormalities. We further show that in the chick embryo, inducing ectopic bone morphogenetic protein (BMP) signaling through in vivo electroporation recapitulates an EA/TEF phenotype. These results reveal both conserved and species-specific characteristics of foregut morphogenesis across mammalian and avian models. The most appropriate disease model therefore depends on the biological scale and mechanism being investigated.

## Results

### The septation-driving mesenchyme and airway patterning are largely conserved in human, mouse, and chick foreguts

The embryonic foregut is patterned along the dorsal-ventral axis into domains that will become the dorsal esophagus and ventral trachea, a process primarily regulated by tissue-scale gradients of BMP, Wnt, and retinoic acid signaling (1, 3, 7). We previously identified a dorsal subepithelial mesenchymal population, labeled with the lineage marker NKX6-1, which we showed to be a critical mechanical driver of epithelial septation through convergent migration in both mouse and chick foreguts (10). In human foregut sections, we found a corresponding NKX6-1-positive cell population in the dorsal subepithelial mesenchyme (Fig. S1A). Expression of FOXF1, a marker of the foregut splanchnic mesoderm lineage, was also highly conserved across species (Fig. S1B).

We next examined dorsal-ventral patterning of the foregut epithelium, which is essential for positioning the tracheal-esophageal boundary in TES. Failure to specify the ventral tracheal domain leads to LTEC (11–13). Ventral expression of the airway epithelial marker NKX2-1 was conserved across human, mouse, and chick foreguts (Fig. S1C). Similarly, phosphorylated SMAD (pSMAD), an indicator of BMP signaling that is critical for specifying airway epithelial identity, showed conserved ventral enrichment across species (Fig. S1D). Expression of the ventral mesenchymal marker ISL1 was also largely conserved (Fig. S1E). Therefore, the major mesenchymal cell populations and dorsal-ventral signaling patterns driving TES are conserved among human, mouse, and chick embryos.

### Mammal-specific patterns of SOX2, SHH, and extracellular matrix

We previously found that SOX2, a marker of esophageal epithelial identity, exhibits contrasting dorsal-ventral expression patterns in mouse and chick foreguts (10). In the chick foregut, SOX2 is expressed broadly throughout the epithelium, with only a modest reduction in the ventralmost tracheal cells (Fig. 1A). By contrast, the mouse foregut exhibits a pronounced dorsal-to-ventral gradient of SOX2 expression (10). We now find that the human foregut, like the mouse, displays a sharp dorsal-to-ventral SOX2 gradient (Fig. 1A and 1B).

**Figure 1.**
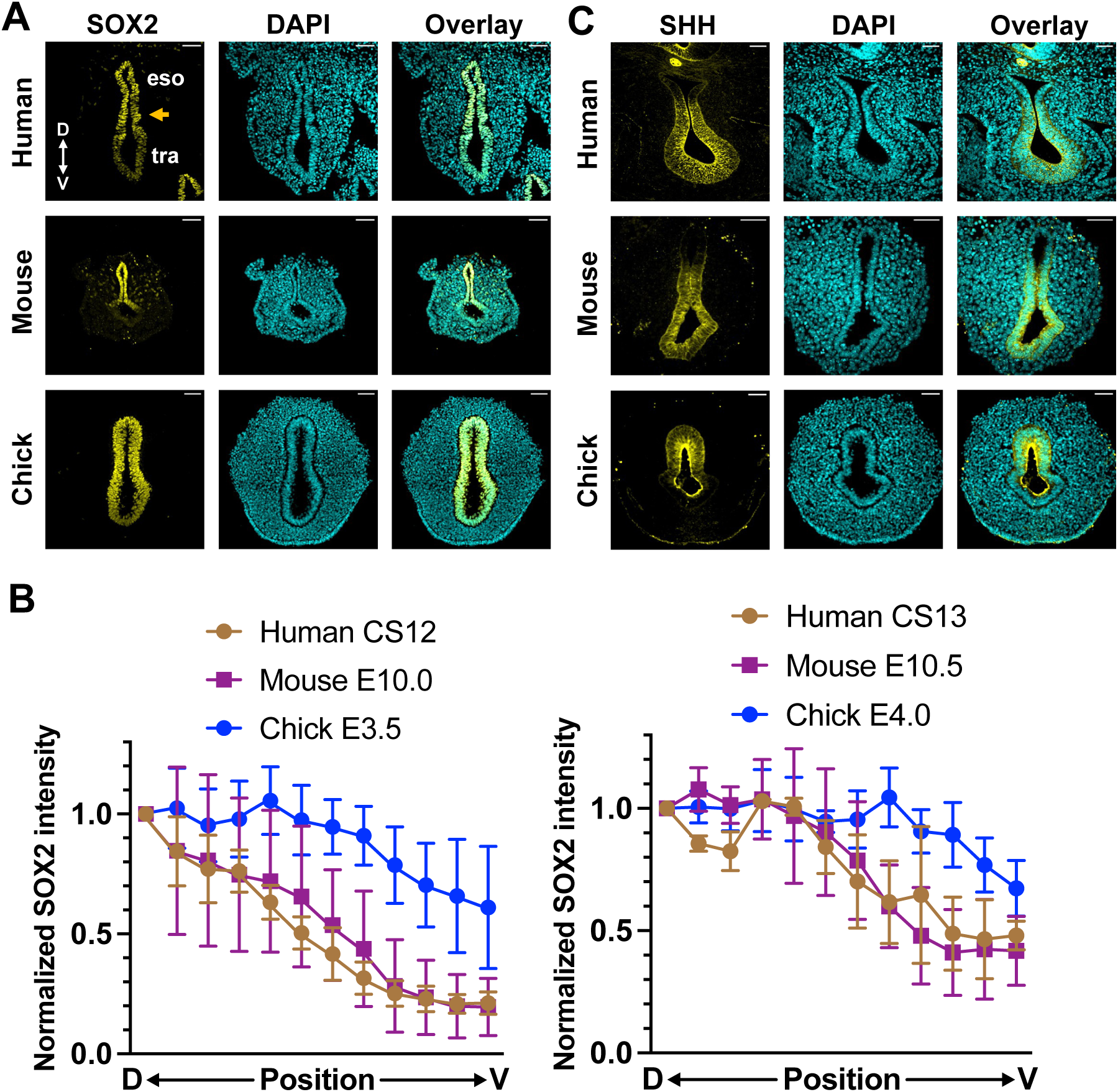
Mammal-specific patterns of SOX2 and SHH. **(A)** Immunofluorescence of SOX2 in human, mouse, and chick foregut sections. The arrow marks the prospective foregut septum. **(B)** Quantification of SOX2 fluorescence along the dorsal-to-ventral axis at different developmental stages. Data are presented as mean ± SD. **(C)** Immunofluorescence of SHH in human, mouse, and chick foregut sections. Images are representative of N = 2 human, N = 4 mouse, and N = 4 chick embryos. D, dorsal; V, ventral; eso, esophagus; tra, trachea. Scale bars, 50 µm.

We next examined Sonic hedgehog (SHH), an important epithelial signal that induces foregut constriction through epithelial-mesenchymal signaling (8, 10, 14). SHH patterning is different between the mouse and chick foreguts (Fig. 1C). In the mouse, SHH is expressed largely throughout the foregut epithelium, with reduced expression in the dorsalmost cells, whereas in the chick it is restricted to the dorsal and medial domains (Fig. 1C). Human SHH expression closely resembles the mouse pattern (Fig. 1C). Expression of the SHH receptor and target gene PTCH1 in the mesenchyme reflects the species-specific distribution of epithelial SHH signaling (Fig. S2A).

A similar interspecies patterning difference is observed for hyaluronic acid (HA), an extracellular matrix component associated with increased tissue fluidity (15). HA is broadly distributed and abundant in the human and mouse foregut mesenchyme, whereas its distribution is more dorsally restricted in the chick (Fig. S2B). These patterns are consistent with our previous live-imaging data, which showed that the mouse mesenchymal cells are broadly dynamic, while in the chick, the dorsal mesenchyme which generates compressive force for TES, is more dynamic than the ventral mesenchyme (10). These results unveil a set of molecular and extracellular matrix patterning features shared by human and mouse foreguts but distinct in the chick, implying mammal-specific gene regulatory mechanisms in the spatial patterning of the foregut.

### Human and chick foreguts share a densely pseudostratified epithelial architecture associated with slower septal resolution

The foregut epithelium is pseudostratified, with densely packed epithelial cells whose nuclei are staggered along the apical-basal axis, while each cell maintains contact with both the apical (luminal) and basal surfaces (16). When we examined epithelial organization across species, to our surprise, we found that the human and chick foregut epithelia appear considerably more densely packed than the mouse foregut (Fig. 2A). We then quantified the line density of epithelial cells in the dorsal (esophageal), medial (prospective septal), and ventral (tracheal) regions of the foregut. The human and chick epithelial cell densities are, respectively, about 1.5 and 2 times that of the mouse epithelium in the dorsal and medial regions (Fig. 2B). Consistent with this difference in epithelial organization, the prospective septum at the onset of TES contained substantially more epithelial cells in the human and chick epithelia than in the mouse (Fig. 2C and 2D).

**Figure 2.**
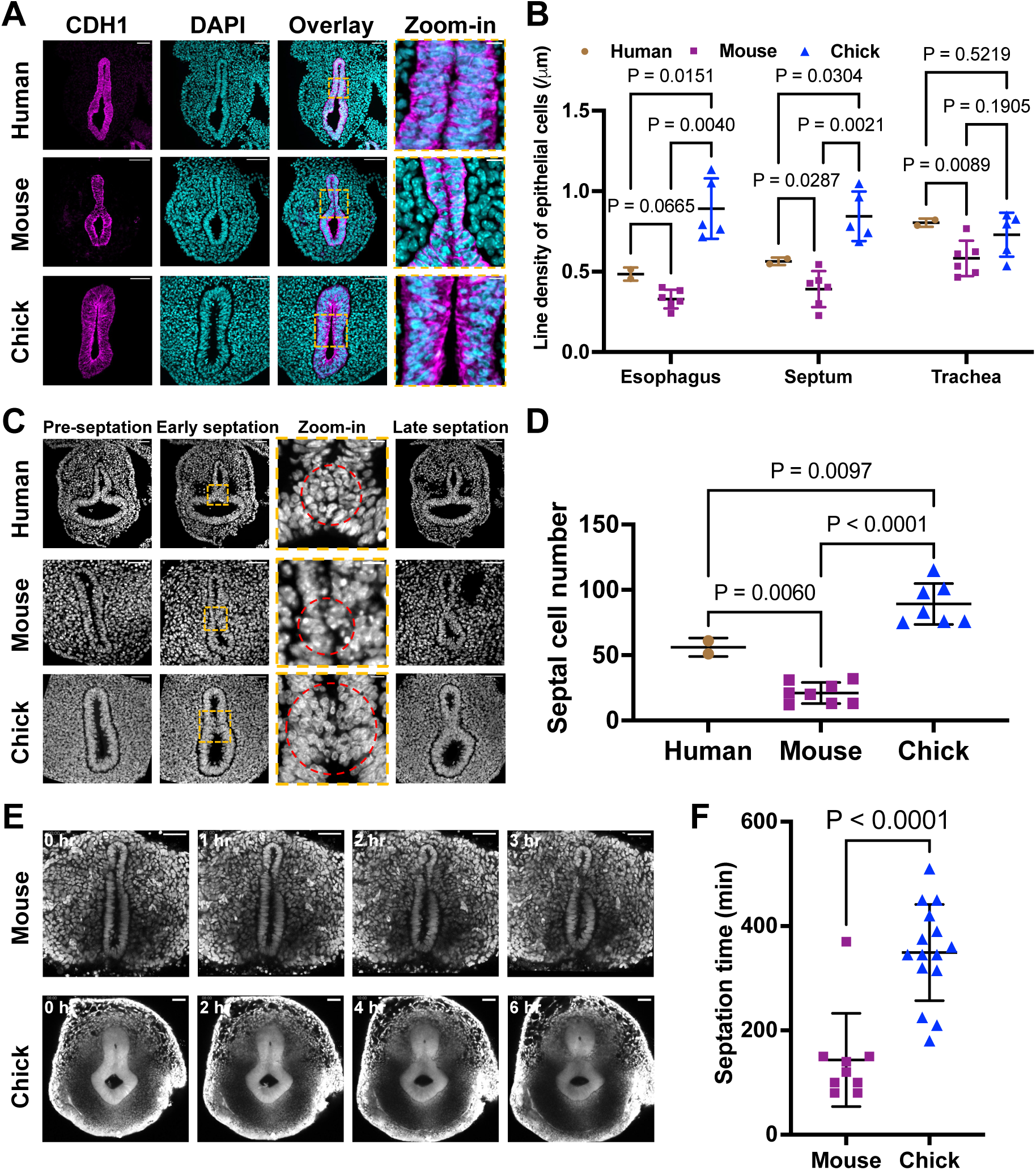
Human and chick foreguts share a densely pseudostratified epithelial architecture associated with slower septal resolution. **(A)** Immunofluorescence of E-cadherin/CDH1 (magenta) in human, mouse, and chick foregut sections, with enlarged views (orange boxes) of the epithelium. **(B)** Line density of epithelial cells in dorsal, medial, and ventral foregut regions. Each point represents one embryo (N = 2 human, 6 mouse, and 5 chick embryos). Data are presented as mean ± SD. P values shown in the graph were calculated by two-way ANOVA with Tukey’s multiple comparisons test. **(C)** Representative DAPI images of morphologically matched pre-septation, early-septation, and late-septation foreguts. Enlarged views (orange boxes) show the epithelial septum. **(D)** Number of cells in the epithelial septum defined in the red ovals in (C). N = 2 human, 8 mouse, and 7 chick embryos. Data are presented as mean ± SD. P values shown in the graph were calculated by one-way ANOVA with Tukey’s multiple comparisons test. **(E)** Two-photon time-lapse imaging of an nTnG mouse foregut slice and a GFP chick foregut slice during septal resolution. Time 0 is the first frame containing a continuous epithelial septum. See also Videos S1 and S2. **(F)** Duration of septal resolution in the mouse (N = 9 embryos) and chick (N = 15 embryos). Data are presented as mean ± SD; P < 0.0001, two-tailed t test. Scale bars, 50 µm; 10 µm in zoom-in views.

These differences in epithelial architecture could influence the dynamics of TES, as septal resolution requires epithelial cells within the prospective septum to remodel their apical-basal polarity (9). Although we did not have access to live human tissue, we compared the duration of septal resolution between the mouse and chick using our previously established slice culture live imaging assay (Fig. 2E, Video S1 and S2) (10). Septal resolution took substantially longer in the chick than in the mouse (Fig. 2F). Together with the similarities in epithelial cell density and septal organization observed in human and chick foreguts, these findings raise the possibility that the dynamics of human septal resolution may resemble the dynamics of the chick.

We then addressed the mechanisms that establish the species-specific differences in foregut epithelial organization. Pseudostratified epithelial architecture is shaped in part by biophysical forces, especially actomyosin-driven apical constriction (16). We previously found that phosphorylated myosin light chain (pMLC), a marker of actomyosin contractility, shows greater apical enrichment in the chick foregut epithelium than in the mouse (10). Unfortunately, pMLC staining was inconsistent between the two human samples, precluding a reliable interspecies comparison (Fig. S3).

To directly test whether actomyosin contractility contributes to the densely packed epithelial architecture observed in the chick foregut, we used azidoblebbistatin, a photoactivatable inhibitor of myosin activity, to selectively reduce actomyosin activity within the epithelium (17). Following patterned two-photon activation of azidoblebbistatin, the chick foregut epithelium gradually became thinner and adopted an organization more similar to that of the mouse (Fig. 3A, Video S3). Prolonged actomyosin inhibition led to extrusion of epithelial cells into the lumen (Fig. 3B, Video S4). Hence, apical constriction is a plausible mechanism that connects the species-specific foregut morphology and TES dynamics.

**Figure 3.**
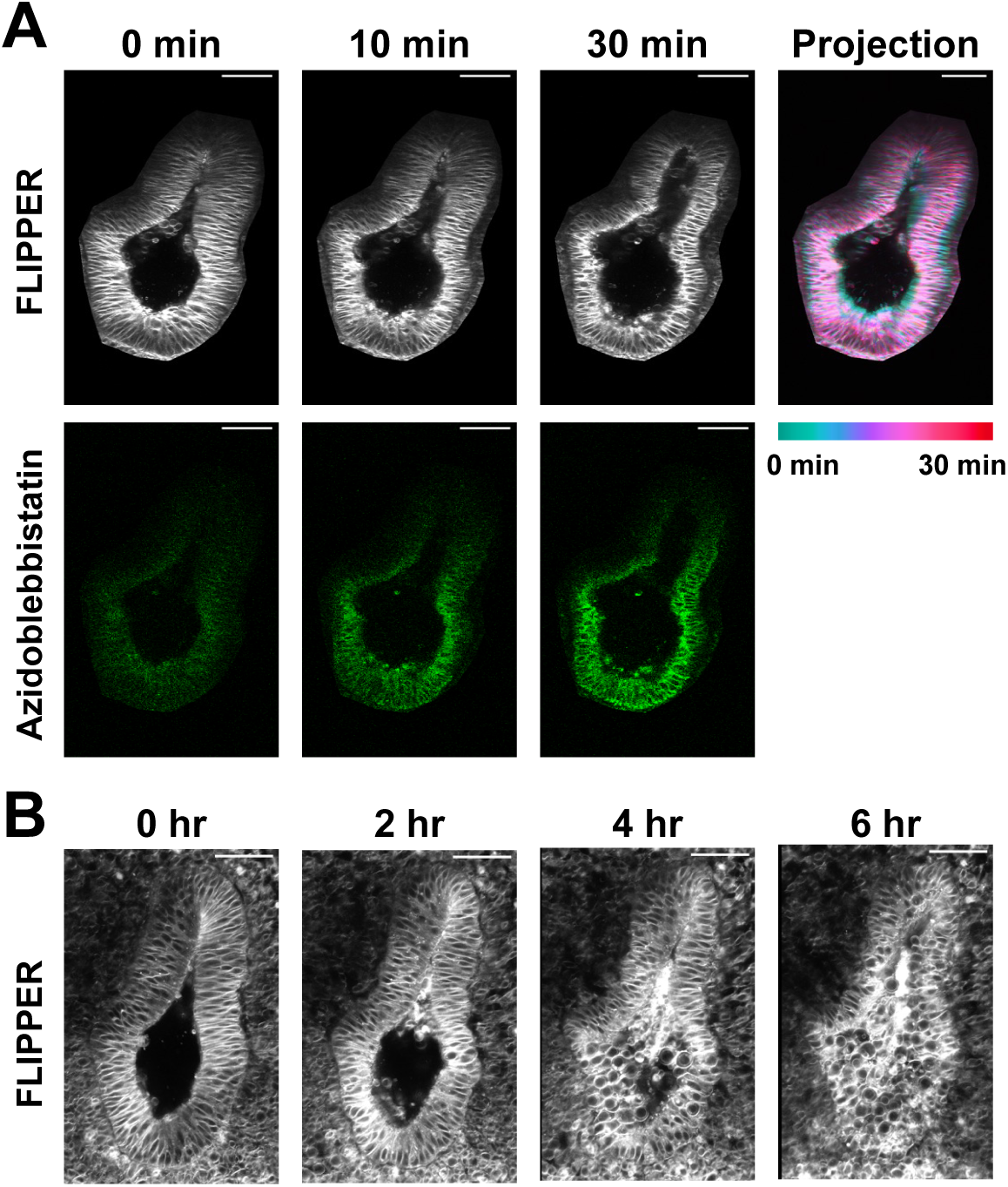
Localized actomyosin inhibition reduces chick foregut epithelial pseudostratification. **(A)** Two-photon live imaging of a FLIPPER-TR-labeled chick foregut slice during epithelial photoactivation of azidoblebbistatin. The temporally color-coded projection summarizes epithelial thinning over the 30-minute recording. See also Video S3. **(B)** Prolonged FLIPPER-TR imaging with continual azidoblebbistatin activation. Continued myosin II inhibition causes progressive epithelial thinning and extrusion of cells into the foregut lumen. Images are representative of N = 4 embryos. See also Video S4. Scale bars, 50 µm.

### Cross-species scRNA-seq analysis reveals conserved foregut cell types and divergent epithelial programs

The similarity between human and chick foregut epithelia prompted us to further investigate whether they also share molecular features different from the mouse. We integrated single-cell RNA-sequencing datasets from CS12-CS14 human embryos (18), E10.5 mouse foreguts (this work), and E3.5-E4.0 chick foreguts (10). Unsupervised clustering resolved conserved cell populations across species (Fig. 4A and S4A), with slight variation in their relative abundance (Fig. 4B and S4B).

**Figure 4.**
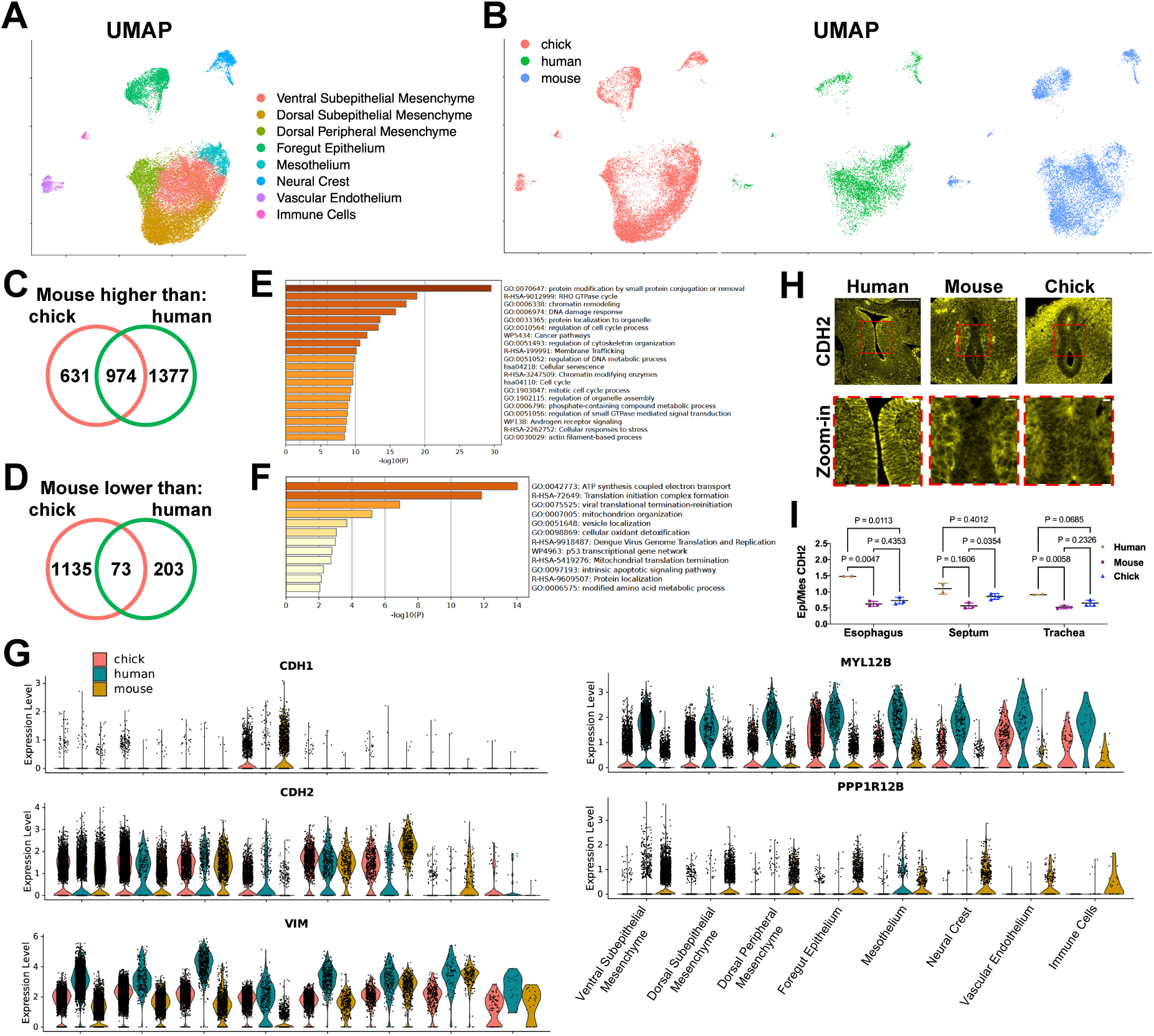
Cross-species scRNA-seq analysis of human, mouse, and chick foreguts. **(A)** UMAP visualization of integrated foregut single-cell transcriptomes from CS12-CS14 human embryos (18), E10.5 mouse foregut slices (this work), and E3.5-E4.0 chick foregut slices (10), colored by Louvain clustering. **(B)** UMAP in (A) split by species. **(C,D)** Differential gene expression analysis of the Foregut Epithelium cluster. Venn diagrams show the overlap between genes expressed at higher (C) or lower (D) levels in the mouse than in the human or chick. Differentially expressed genes were defined by an absolute log2FC>1 and adjusted P < 0.001. **(E,F)** Metascape functional enrichment analysis of the overlapping genes expressed at higher (E) or lower (F) levels in the mouse than in the human and chick. **(G)** Violin plots of expression levels of CDH1 (E-cadherin), CDH2 (N-cadherin), VIM (vimentin), MYL12B (MLC), and PPP1R12B (MYPT2) in all cell clusters split by species. **(H)** Immunofluorescence of N-cadherin/CDH2 in human, mouse, and chick foregut sections, with enlarged views (red boxes) highlighting the prospective septal region. (I) Quantification of epithelial-to-mesenchymal N-cadherin intensity ratios in dorsal, medial, and ventral foregut regions. Each point represents one embryo (N = 2 human, 3 mouse, and 3 chick embryos). Data are presented as mean ± SD. P values shown in the graph were calculated by two-way ANOVA with Tukey’s multiple comparisons test. Scale bars, 50 µm in overviews, and 10 µm in zoom-in views.

We next compared key patterning genes within the dorsal subepithelial mesenchyme, which generates the compressive force driving TES (10). We found that Wnt-, ephrin-, and semaphorin-signaling genes show strong mammal-chick divergence, whereas SHH-response genes were expressed at similar levels across all species (Fig. S4C).

Within the foregut epithelium, we performed pairwise interspecies gene expression comparisons, and identified genes whose expression levels were higher or lower in the mouse than in the human or chick (Fig. 4C and 4D). Functional enrichment of the common mouse-high genes revealed processes associated with RHO GTPases, cytoskeletal organization, and cell cycle (Fig. 4E). Conversely, the common mouse-low genes were enriched for biosynthetic metabolism (Fig. 4F). These results suggest that interspecies epithelial divergence in morphology involves multiple regulatory and metabolic programs.

Focusing on genes associated with epithelial adhesion and actomyosin regulation, we found that E-cadherin/CDH1 expression was higher, whereas N-cadherin/CDH2 and vimentin were lower in the mouse epithelium compared to human and chick epithelia (Fig. 4G). MYL12B, encoding a myosin light chain (MLC), was expressed at lower levels in the mouse than in the human or chick (Fig. 4G). In contrast, PPP1R12B, encoding the myosin phosphatase-targeting subunit MYPT2, was upregulated in the mouse (Fig. 4G). These expression patterns are consistent with species-specific regulation of actomyosin activity and with the less pseudostratified epithelial architecture observed in the mouse foregut.

The difference in N-cadherin is particularly intriguing. Although N-cadherin is conventionally associated with the neural epithelium and mesenchymal tissues, we found considerable CDH2 transcripts in the foregut epithelium in all three species, with the mouse foregut expressing it at a significantly lower level (Fig. 4G). We validated N-cadherin expression using immunofluorescence. N-cadherin was observed in the mesenchyme in all three species but showed variable enrichment within the epithelium, particularly in the prospective septal region (Fig. 4H). Quantification of epithelial-to-mesenchymal fluorescence ratios showed greater epithelial enrichment in the human and chick foreguts than the mouse, especially in the septal region (Fig. 4I). The co-expression of E-cadherin, N-cadherin, and vimentin may reflect a more plastic epithelial state primed for polarity remodeling during TES. Intriguingly, previous observation in the mouse intestine, in which substitution of E-cadherin with N-cadherin promotes epithelial hyperplasia and polyp formation, suggests that changes in cadherin composition can influence epithelial organization (19).

Together, these findings suggest that species-specific regulation of cell adhesion and actomyosin activity may contribute to the convergence of human and chick epithelial architecture and its divergence from the mouse.

### Modeling human EA/TEF in the chick embryo

Given the structural similarities between human and chick foregut epithelia, we next asked whether genetic perturbation in the chick embryo would recapitulate human EA/TEF phenotypes. Clinical and genetic studies have implicated elevated BMP signaling in the pathogenesis of EA/TEF, owing to reduced expression of the BMP antagonist NOGGIN in the dorsal foregut (20). The Noggin-knockout mouse is one of the few models that reproduces this human malformation, albeit with incomplete penetrance (1, 21, 22). We thus hypothesized that ectopic elevation of BMP signaling in the dorsal chick foregut would similarly disrupt TES. We overexpressed BMP7 in the dorsal foregut epithelium through in vivo electroporation. Compared with control embryos (Fig. 5A), BMP7 overexpression caused marked hypoplasia and discontinuity of the esophagus, generating an EA/TEF-like foregut configuration (Fig. 5B).

**Figure 5.**
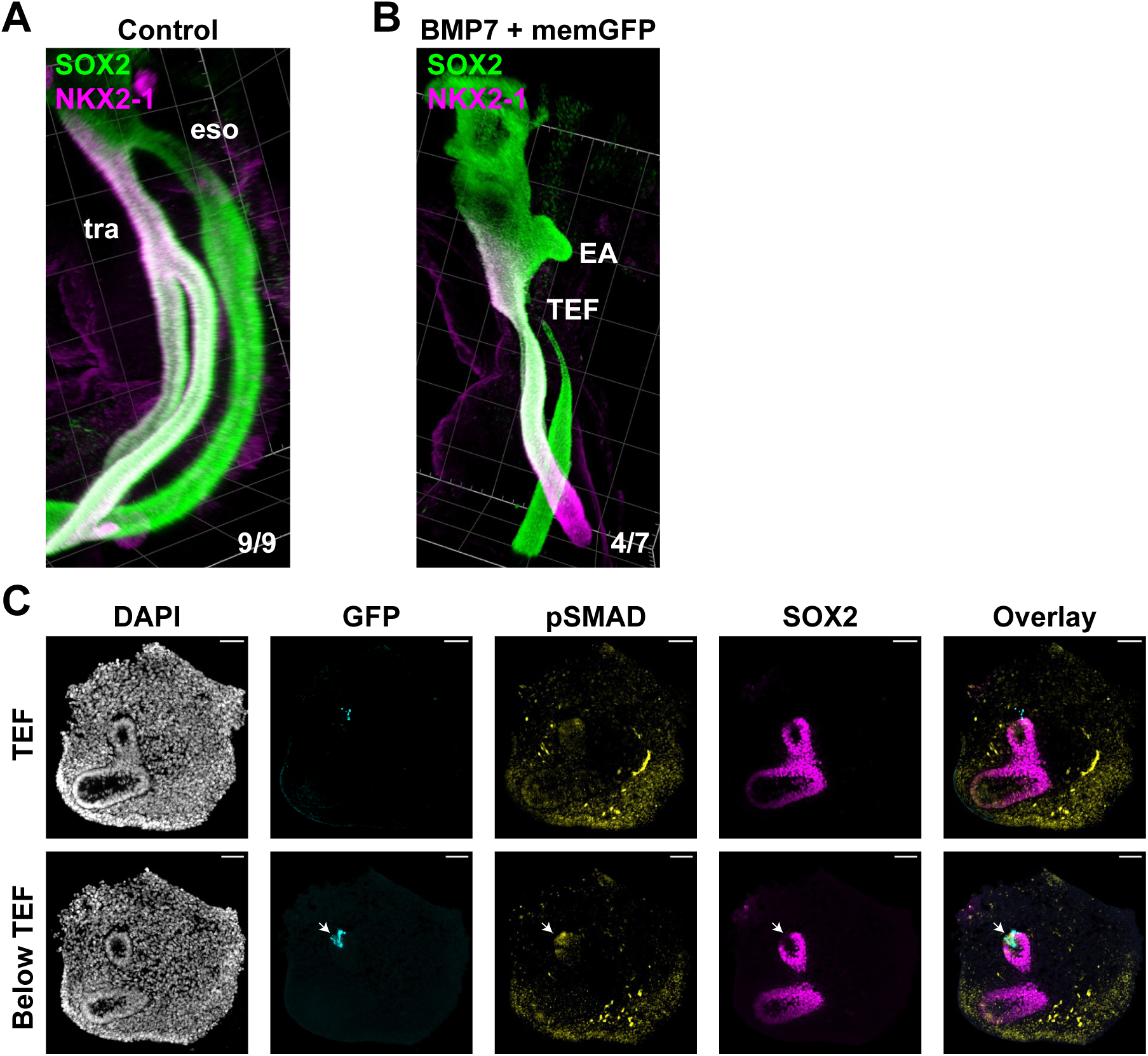
Epithelial BMP7 overexpression produces an EA/TEF-like phenotype in chick embryos. **(A)** Whole mount immunofluorescence of SOX2 (green) and NKX2-1 (magenta) in a control E5 chick foregut. All control embryos completed tracheal-esophageal separation (9/9). **(B)** Whole mount immunofluorescence of SOX2 and NKX2-1 after co-electroporation of BMP7 and membrane-GFP. Four of seven embryos developed esophageal hypoplasia with an ectopic tracheoesophageal fistula (TEF). **(C)** Transverse sections through the TEF and at a level below the TEF. BMP7-expressing epithelium shows reduced SOX2 and induces ectopic pSMAD in adjacent tissue (arrow). Images are representative of N = 3 electroporated embryos. Grid size, 200 µm (A, B). Scale bars, 50 µm (C).

Sections through the TEF revealed substantial changes in foregut morphology and dorsal-ventral patterning (Fig. 5C). Locally elevated BMP7 reduced SOX2 expression in the dorsal epithelium and induced ectopic BMP signaling in the adjacent mesenchyme (Fig. 5C), consistent with the Noggin-knockout mouse embryo phenotype (21). These results demonstrate that targeted perturbation of BMP signaling in the chick foregut can reproduce key anatomical and molecular features of EA/TEF, providing proof of principle for using chick embryos as a model for studying human foregut malformations, particularly those involving epithelial abnormalities.

## Discussion

Motivated by the discrepancies between mouse and chick TES in molecular patterning and tissue dynamics, our study unexpectedly reveals mixed characteristics of the human foregut. The human foregut resembles the mouse in its dorsal-ventral molecular patterning but resembles the chick in its epithelial architecture, exhibiting substantially greater pseudostratification than the mouse. The extent of epithelial pseudostratification correlates with the dynamics of septal resolution, implying that septum resolution during human TES may proceed through more chick-like epithelial dynamics. Thus, molecular patterning and epithelial morphogenesis can vary independently across species, and neither mouse nor chick alone fully captures human TES.

The developmental origin of this divergence in epithelial architecture remains an open question. One possibility is that it reflects differences in the morphogenetic history of the foregut. During gastrulation, human and chick embryos initially develop as flat discs of three germ layers, whereas the mouse embryo has a cup-shaped, cylindrical geometry (23). These distinct embryonic structures impose different biophysical constraints during foregut formation. In the chick, actomyosin-driven apical constriction plays an essential role in folding the endoderm into the foregut tube (24), whereas the mouse foregut is formed through endodermal involution independent of actomyosin activity (25, 26). These potentially divergent biomechanical drivers of foregut formation may underlie the differences in epithelial organization. Consistently, the chick foregut exhibits greater apical actomyosin enrichment than the mouse (10), and local inhibition of actomyosin with photoactivated azidoblebbistatin reduces epithelial pseudostratification. Consistent with these findings, cross-species scRNA-seq analysis reveals tissue-level downregulation of MLC and upregulation of MYPT2 in the mouse foregut relative to the human and chick, as well as epithelium-specific downregulation of N-cadherin, which potentially contributes to pseudostratification given its high expression in the highly pseudostratified neural tube (27). Notably, substituting E-cadherin with N-cadherin in the mouse intestine leads to overgrowth features including polyp formation (19).

The differences in epithelial organization may also influence susceptibility to distinct congenital malformations. EA/TEF accounts for more than 70% of human TES defects, yet it is reproduced in only a small number of mouse models, such as the Noggin-knockout and Sox2-hypomorphic models (22, 28). By contrast, more than 20 mouse models of defective TES predominantly generate the LTEC phenotype with a completely unseparated foregut (3, 4). As LTEC is mainly driven by the defects of mesenchymal cells to initiate epithelial constriction through convergent migration (8, 10), EA/TEF may instead result from a later or partial failure of TES, in which constriction begins but epithelial remodeling and separation are not coordinated correctly (9).

Compared with the relatively simple, rapidly resolving mouse foregut epithelium, the more highly pseudostratified epithelia of human and chick foreguts may require prolonged and more coordinated epithelial remodeling during septal resolution, making them more susceptible to incomplete separation such as EA/TEF.

Our work establishes the chick as a complementary model for investigating the epithelial basis of human foregut malformations. With the ability to introduce spatially and temporally controlled perturbations through in vivo electroporation, and to monitor TES morphogenesis through explant culture and live imaging, the chick embryo provides a tractable system for dissecting how genetic perturbations in the foregut epithelium lead to congenital malformations. More broadly, our results argue that experimental models should be selected according to the specific molecular or tissue-level mechanism under investigation.

## Acknowledgements

We thank Nita Solanky, Steven Lisgo, and Jacqui Dobor of the Human Developmental Biology Resource for providing the fixed human embryonic samples. We thank Suzanne White at the BIDMC Histology Core for human sample processing, Susan Chapman for providing GFP chick eggs, the MicRoN Facility of Harvard Medical School for microscopy resources, and the Center for Comparative Medicine at Harvard Medical School for mouse colony maintenance. This work was supported by a Helen Hay Whitney Fellowship to R.Y. and a discretionary fund made available to C.J.T. by Harvard Medical School.

## Author contributions

R.Y. conceived the project, performed the experiments, and analyzed the data. A.Z.M. analyzed the scRNA-seq datasets with R.Y. C.J.T. supervised the project. R.Y. and C.J.T. wrote the manuscript with A.Z.M.

## Methods

All animal studies were performed in compliance with the protocols approved by the Institutional Animal Care and Use Committee at Harvard Medical School.

### Human embryonic samples

Fixed human embryonic samples were provided by the Human Developmental Biology Resource (HDBR). Tissue was collected after elective termination of pregnancy with written informed maternal consent under approvals held by HDBR. Samples were de-identified before transfer. HDBR assigned developmental stages from external morphology and measurements according to Carnegie staging. We processed one Carnegie Stage 12 (CS12) embryo and one CS13 embryo.

### Mouse embryos

C57BL/6J (The Jackson Laboratory, 000664) and ROSA-nT-nG (nTnG; The Jackson Laboratory, 023537) mice were used. Animals were maintained on a 12-hour light/dark cycle with food and water available ad libitum. Timed pregnancies were set up, and the day of detecting a vaginal plug was designated as embryonic day (E) 0.5. Embryos were collected between E9.5 and E10.5.

### Chick embryos

Fertilized White Leghorn eggs were obtained from Charles River Laboratories (AVS Bio). Roslin Green transgenic eggs, which constitutively express untagged GFP, were provided by Susan Chapman. Eggs were incubated at 38°C in a humidified chamber. Embryos from E3.5-E4.5 were used.

### Foregut slice culture

Fresh chick or mouse foregut slice culture was performed as described (10). Briefly, the foregut from the pharyngeal arches to the upper stomach was isolated and cut transversely into 100-150 µm slices with a surgical blade (Aspen Surgical, 371111). Slices containing the tracheal-esophageal septum were embedded in Matrigel (Corning, 356231). Culture medium was added after gelation at 37°C for 30 minutes. Slices were maintained at 37°C with 5% CO₂ until imaging.

### Two-photon time-lapse imaging

Live imaging was performed on a Leica Stellaris 8 multiphoton microscope equipped with an Insight X3 dual-beam laser and a 25X/1.0 NA water-immersion objective (Leica, 11507703). GFP chick slices were excited at 960 nm with 7% laser power, and nTnG mouse slices were excited at 1040 nm with 7% laser power. For each slice, a 30-60 µm z-stack was acquired with a 10-15 µm step size, beginning approximately 30 µm below the cut surface. Samples were recorded every 5-15 minutes for 12-16 hours at 250-500 nm per pixel in the xy plane. The duration of septal resolution was measured from the first frame when a continuous epithelial septum bridged the foregut lumen to the first frame when the junction had completely resolved.

### Two-photon photoactivation of azidoblebbistatin

Chick foregut slices were labeled with 2 µM FLIPPER-TR (Cytoskeleton, CY-SC020) in Dissection Medium for 30 minutes at room temperature with intermittent agitation. Labeled slices were embedded in Matrigel and cultured as described above in medium containing 0.5 µM azidoblebbistatin (Motorpharma). FLIPPER-TR was excited at 1040 nm with 7% laser power to monitor tissue morphology without activating azidoblebbistatin. Azidoblebbistatin was locally photoactivated at 860 nm with 10-15% laser power in a user-defined region of interest encompassing the foregut epithelium. Samples were recorded every 12 seconds for 30 minutes, and longer recordings continued for 6 hours to assess tissue integrity and luminal cell extrusion at 10 minutes per frame.

### In vivo electroporation of the chick foregut epithelium

pCAG-BMP7 (Addgene 163591) and pCAGGS-myrGFP (Addgene 115502) were co-electroporated at a 4:1 ratio. The electroporation mixture contained 4 µg/µL plasmid DNA, 0.5% Fast Green FCF (Sigma-Aldrich, F7252), and 3% sucrose (Sigma-Aldrich, S8501) in TE buffer (Qiagen, 19086). E2 chick embryos were lowered and windowed, and approximately 30 µL of 1:20 diluted ink (Pelikan, 211862) in PBS was injected into the yolk beneath the embryo for visualization. A pulled glass capillary (FHC, 30-30-0; Sutter Instrument P-97 puller) was used to inject plasmid solution into the foregut lumen until the dye reached the anterior intestinal portal. A parallel needle electrode (Bulldog Bio, CUY560-5-0.5) was inserted into the yolk along the anterior-posterior axis, with the positive electrode on the right side. Three 50-V, 7-ms poring pulses separated by 100 ms were followed by five 20-V, 7-ms transfer pulses separated by 100 ms using a NEPA21 electroporator (Nepa Gene). Fresh albumen was added over the embryo. The eggs were sealed and returned to the incubator until collection at E5.

### Whole mount immunofluorescence

Whole mount immunofluorescence was performed as described (10). Fixed foreguts were permeabilized, blocked, and incubated with primary antibodies for 2 days at 4°C, followed by three washes with the blocking buffer at room temperature, and incubation with fluorescent secondary antibodies for 1 day at 4°C. Samples were extensively washed, serially dehydrated into methanol, and optically cleared with the CytoVista Tissue Clearing kit (Invitrogen, V11322). Cleared specimens were imaged on a Nikon Ti inverted microscope with a W1 spinning-disk scanner (Yokogawa CSU-W1) using a 10× objective and 0.6-1.0 µm z-steps. Three-dimensional renderings were made in Arivis Vision4D (Zeiss), with the z-step multiplied by 1.5 to correct for refractive-index mismatch (29).

### Immunofluorescence of mouse and chick tissue sections

Mouse and chick embryos, dissected foreguts, and cultured slices were fixed overnight at 4°C in 4% paraformaldehyde in PBS. Samples were incubated sequentially in 15% sucrose, 30% sucrose, and 1:1 O.C.T. Compound (Sakura Finetek, 4583):30% sucrose for 1 hour per step at room temperature, embedded in O.C.T., frozen in a dry ice-ethanol bath, and stored at -80°C. Transverse sections at 14 µm thickness were collected on Superfrost Plus slides (Fisher Scientific, 12-550-15) using a Leica CM3050 cryostat. Slides were dried for 20 minutes at 50°C, and rinsed with PBS for three times to remove O.C.T. For SHH and phospho-myosin light chain staining, antigen retrieval was performed for 10 minutes in boiling citrate buffer (Abcam, ab64214). Sections were permeabilized for 20 minutes with 0.1% Triton X-100 in PBS and blocked for 1 hour in 5% donkey serum (Jackson ImmunoResearch, 017-000-121) and 0.3% Triton X-100 in PBS. Primary antibodies were applied overnight at 4°C. Sections were washed three times for 10 minutes in blocking buffer diluted 1:10 in PBS, incubated for 2 hours at room temperature with fluorescent secondary antibodies and 10 µg/mL DAPI (Invitrogen, D1306), washed three times, and mounted in ProLong Diamond Antifade Mountant (Invitrogen, P36970).

### Immunofluorescence of human tissue sections

Formalin-fixed, paraffin-embedded (FFPE) human embryo samples were remounted in paraffin to cut transverse sections. Serial sections of 5 µm thickness were prepared by the BIDMC Histology Core. Sections were deparaffinized by serial 3-minute washes in xylene (Fisher Scientific, X5-4, twice), 100% ethanol (Koptec, V1016, twice), 95% ethanol, 70% ethanol, 50% ethanol, and water (three times). After three PBS washes, antigen retrieval was performed for 10 minutes in boiling HistoVT One buffer (Nacalai, 06380-76). Samples were then immunostained as described above.

### Antibodies

Primary antibodies and probes were rat anti-SOX2 (1:300; Invitrogen, 14-9811-82), rabbit anti-NKX2-1 (1:300; Abcam, ab76013), goat anti-NKX6-1 (1:100; R&D Systems, AF5857), goat anti-FOXF1 (1:100; R&D Systems, AF4798), mouse anti-ISL1 (1:100; Developmental Studies Hybridoma Bank, 40.2D6), goat anti-SHH (1:100; R&D Systems, AF464), rabbit anti-phospho-SMAD1/5/9 (1:100; Cell Signaling Technology, 13820), mouse anti-CDH1 (1:150; BD Biosciences, 610182), mouse anti-CDH2 (1:100; BD Biosciences, 610920), rabbit anti-phospho-myosin light chain 2 (1:100; Cell Signaling Technology, 3674), chick anti-GFP (1:500; Abcam, ab13970), and biotinylated hyaluronic acid-binding protein (1:200; Sigma-Aldrich, 385911). Alexa Fluor 488-, Cy3-, or Alexa Fluor 647-conjugated secondary antibodies (Jackson ImmunoResearch) were used at 1:300. Hyaluronic acid-binding protein was detected with streptavidin-Alexa Fluor 647 (1:300; Invitrogen, S32357).

### Hybridization chain reaction (HCR)

HCR v3.0 probes targeting human PTCH1, mouse Ptch1, and chick PTCH1 were designed from their respective coding sequences using insitu_probe_generator (https://github.com/rwnull/insitu_probe_generator). Candidate probes were searched against the corresponding transcriptome to remove predicted off-target sequences, appended with Molecular Instruments HCR initiators, synthesized as 50-pmol oPools (Integrated DNA Technologies), and reconstituted to 1 µM in TE buffer. Mouse and chick foreguts were stained as whole mounts before being embedded and sectioned, while human samples were stained on sections. Samples were permeabilized in 70% ethanol for 1 hour at room temperature, equilibrated for 10 minutes in HCR probe wash buffer and for 30 minutes at 37°C in hybridization buffer (Molecular Instruments), and hybridized overnight at 37°C with 40 nM probe. Samples were washed twice in probe wash buffer, twice in 5X SSCT (5X saline-sodium citrate [Invitrogen, 15557044] and 0.1% Tween 20 [Sigma-Aldrich, P9416]), and once in amplification buffer for 20 minutes per wash. Alexa Fluor 546-conjugated hairpins were denatured for 90 seconds at 95°C, annealed at room temperature, diluted to 30 nM per strand in amplification buffer, and incubated with samples overnight at room temperature. Samples were finally washed twice in 5X SSCT and once in PBS.

### Microscopy and image analysis

Stained tissue sections were imaged on a Nikon Ti inverted microscope equipped with a W1 spinning-disk scanner (Yokogawa CSU-W1) using 20X and 40X objectives. Z-stacks were acquired at 0.3-0.9 µm intervals. A single z section was shown in Fig. 2 and Fig. 3, while in other figures images were displayed as maximum intensity projections. Images compared within an experiment were acquired and processed with the same settings.

Dorsal-ventral fluorescence profiles were quantified in Fiji as described (10). For epithelial line density, DAPI-positive epithelial nuclei were counted within manually delineated dorsal esophageal, medial septal, and ventral tracheal segments and divided by the corresponding basal epithelial contour length. Septal cell number was determined by manual counting of DAPI-labeled nuclei within the epithelial junction at the onset of septation. Septation duration was calculated from live recordings as described above. N-cadherin intensity was determined by manually drawing polygonal lines along the cell boundary in Fiji, and the measured intensity was subtracted by the background intensity measured inside the foregut lumen. Multiple sections from the same embryo were averaged as one biological replicate.

### Mouse foregut single-cell RNA sequencing (scRNA-seq)

Sample preparation, library construction, and sequencing of E10.5 mouse foregut slices were performed in the same batch as the previously reported chick foreguts (10). Briefly, control E10.5 mouse embryos (N = 9) were dissected to isolate the foreguts, which were sliced to enrich the regions undergoing TES. Slices were dissociated in TrypLE (Gibco, 12604013) at 37 °C. To distinguish the mouse sample (9 embryos pooled together) from the chick, we performed the MULTI-seq workflow (30). Library construction was performed with the Chromium Next GEM Single Cell 3’ Kit (v3.1) with dual index (10x Genomics). Quality control of the library was performed by the Biopolymers Facility at Harvard Medical School. The 10-nM library was sequenced with the NovaSeq 6000 platform at the Biopolymers Facility at Harvard Medical School.

### Cross-species integration and analysis of scRNA-seq datasets

Preprocessed single-cell RNA-sequencing datasets from human, mouse, and chick foreguts were integrated in R using Seurat v5.0.1 (31). To harmonize genes across species, chick and mouse genes were converted to their human ortholog symbols using the Ensembl BioMart database (release 116) with the biomaRt package. Only ortholog pairs classified by Ensembl as one-to-one were retained. Following orthology conversion, all three datasets were trimmed to the 10,166 unique human genes. As the human dataset was from whole embryos across stages (18), we subset only CS12-CS14 foregut cells based on known markers. The resulting dataset contained 24,634 cells, comprising 4,200 human, 7,890 mouse, and 12,544 chick cells.

After cell cycle regression, we performed principal-component analysis using 2,000 variable genes, and the first 30 principal components were used for cross-species integration by canonical correlation analysis with the Seurat v5 IntegrateLayers and CCAIntegration functions. Cells were grouped by graph-based Louvain clustering at a resolution of 0.2, producing 8 clusters. Uniform Manifold Approximation and Projection (UMAP) was used to visualize cells by cluster and species. The clusters were annotated according to known markers of the foregut cell types (10). Ventral Subepithelial Mesenchyme was identified by ISL1, Dorsal Subepithelial Mesenchyme by PDE1A, Dorsal Peripheral Mesenchyme by TBX1, Foregut Epithelium by EPCAM, Mesothelium by ALDH1A2, Neural Crest by SOX10, Vascular Endothelium by PECAM1, and Immune Cells by CSF1R. To compare cell-type composition across species, the number of cells assigned to each annotated cluster was divided by the total number of cells from the corresponding species.

Differential gene expression analysis was performed with the FindMarkers function in a pairwise fashion to avoid interspecies averaging. Genes were classified as differentially expressed based on adjusted P <0.001 and absolute log2FC >1. Genes expressed at higher or lower levels in the mouse than in both the human and chick were identified by intersecting the mouse-versus-human and mouse-versus-chick differential gene sets. The shared mouse-high and mouse-low gene sets were analyzed separately using Metascape (32). The most significantly enriched terms were ranked and plotted according to their P values.

## Statistics and reproducibility

Sample sizes and statistical tests are indicated in the figure legends. The sample sizes were not predetermined. GraphPad Prism was used to plot the data.

## Data availability

The human embryo scRNA-seq dataset was from (18) with Gene Expression Omnibus (GEO) accession code GSE157329. The chick foregut scRNA-seq dataset was from (10) with GEO accession code GSE307539. The mouse foregut scRNA-seq data and the integrated, cross-species dataset will be uploaded to the GEO database.

## Code availability

No custom code was generated in this study.

---

**Figure S1.**
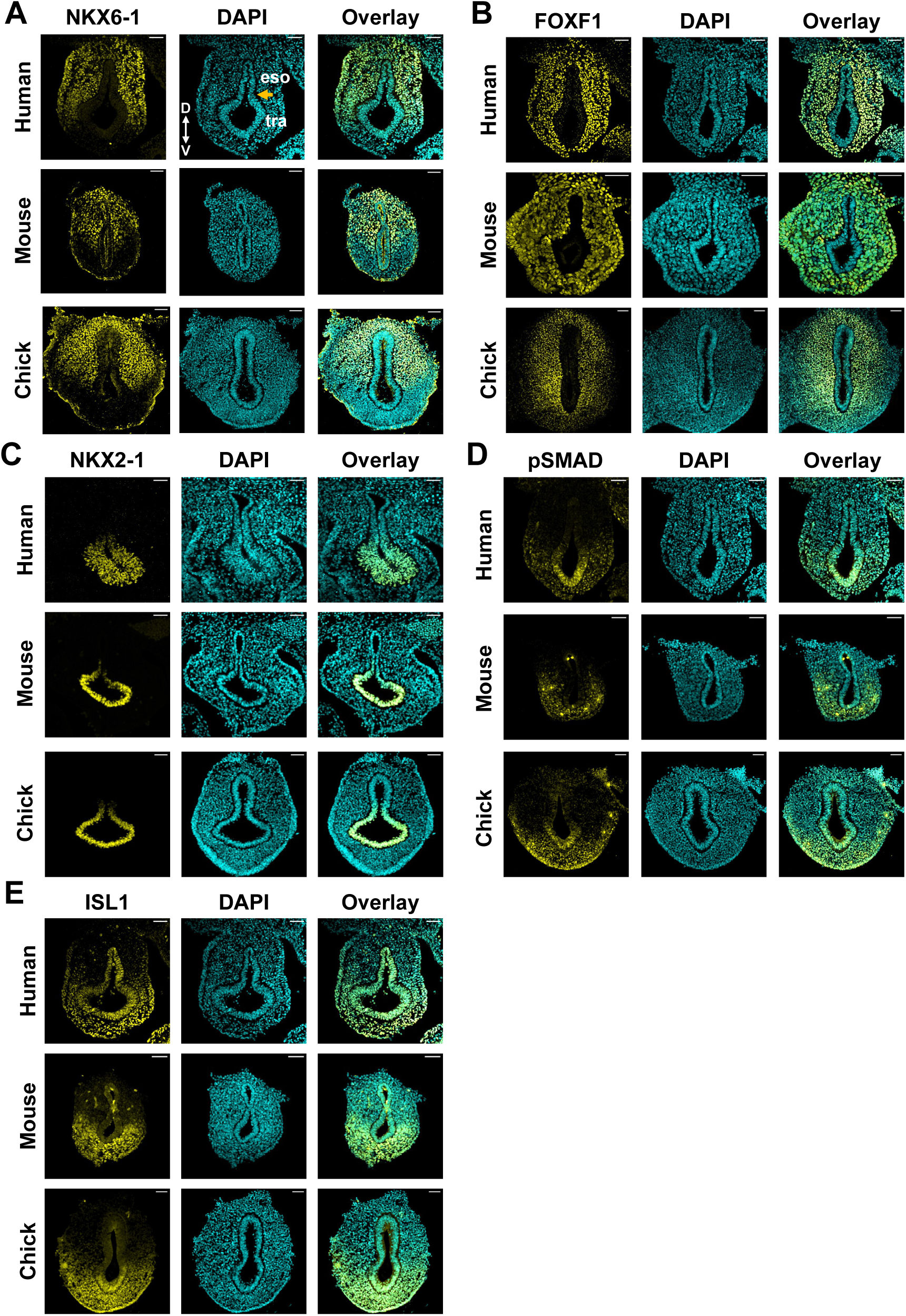
The septation-driving mesenchyme and airway patterning are largely conserved in human, mouse, and chick foreguts. **(A)** Staining of NKX6-1 (yellow) and DAPI (cyan) in transverse foregut sections at developmentally matched stages of tracheal-esophageal separation. **(B)** Immunofluorescence of the pan-foregut mesenchyme marker FOXF1 in transverse foregut sections. **(C)** Immunofluorescence of NKX2-1 in human, mouse, and chick foreguts. **(D)** Immunofluorescence of phospho-SMAD1/5/9 (pSMAD; yellow) in human, mouse, and chick foreguts. **(E)** Immunofluorescence of ISL1, showing enrichment in ventral foregut mesenchyme across species. Images are representative of N = 2 human, N = 4 mouse, and N = 4 chick embryos. D, dorsal; V, ventral; eso, esophagus; tra, trachea. Scale bars, 50 µm.

**Figure S2.**
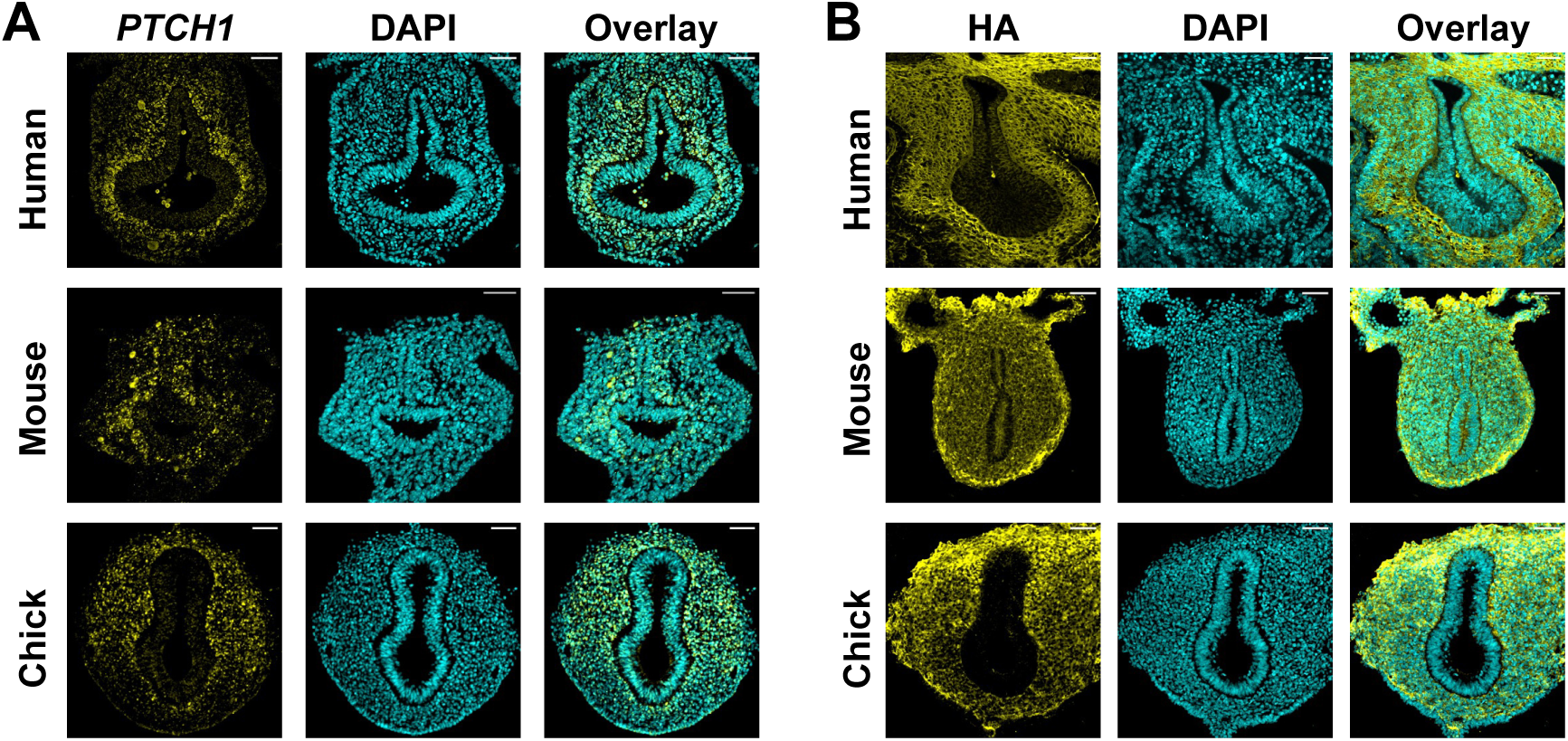
PTCH1 and hyaluronic acid distributions in human, mouse, and chick foreguts. **(A)** HCR fluorescence in situ hybridization for PTCH1 transcripts in transverse foregut sections. PTCH1 expression reflects the broader SHH-responsive mesenchyme in the human and mouse and a dorsally biased domain in the chick. **(B)** Hyaluronic acid detected with biotinylated hyaluronic acid-binding protein. Human and mouse foreguts show a broader HA distribution, whereas chick HA is enriched dorsally. Images are representative of N = 2 human, N = 4 mouse, and N = 4 chick embryos. Scale bars, 50 µm.

**Figure S3.**
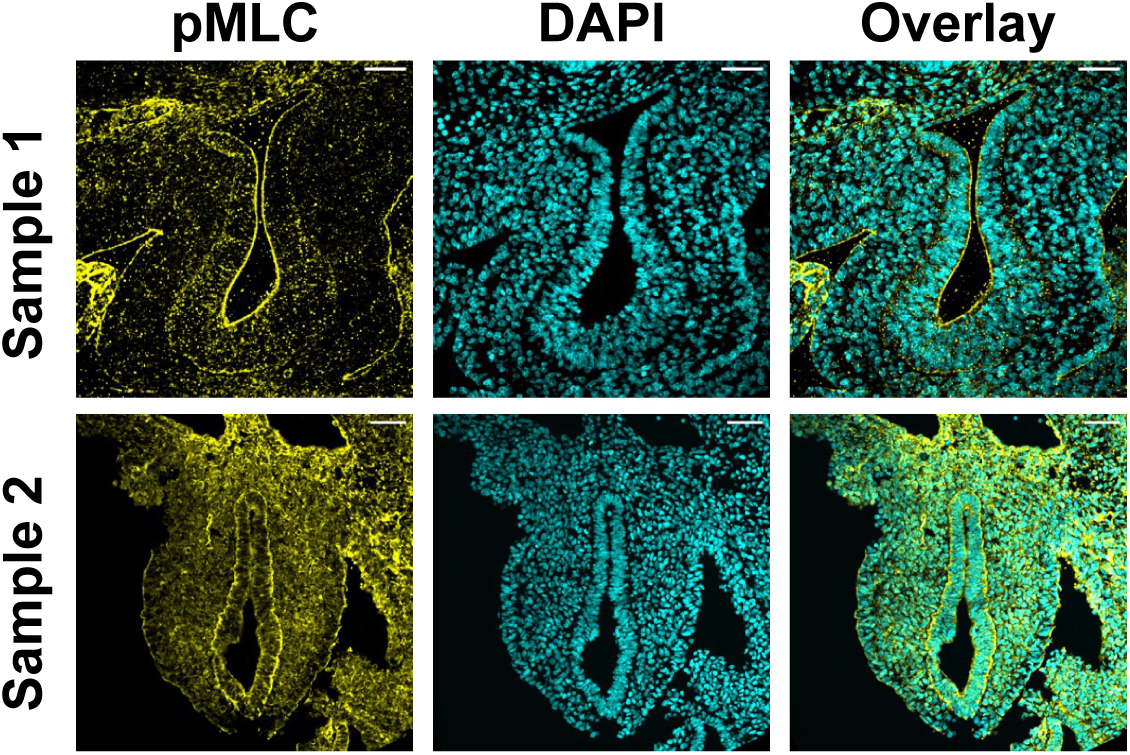
Phosphorylated myosin light chain staining in two human foregut samples. Immunofluorescence of phospho-myosin light chain 2 (pMLC) in transverse sections from two independent human embryonic samples. The samples show variable apical pMLC enrichment. Scale bars, 50 µm.

**Figure S4.**
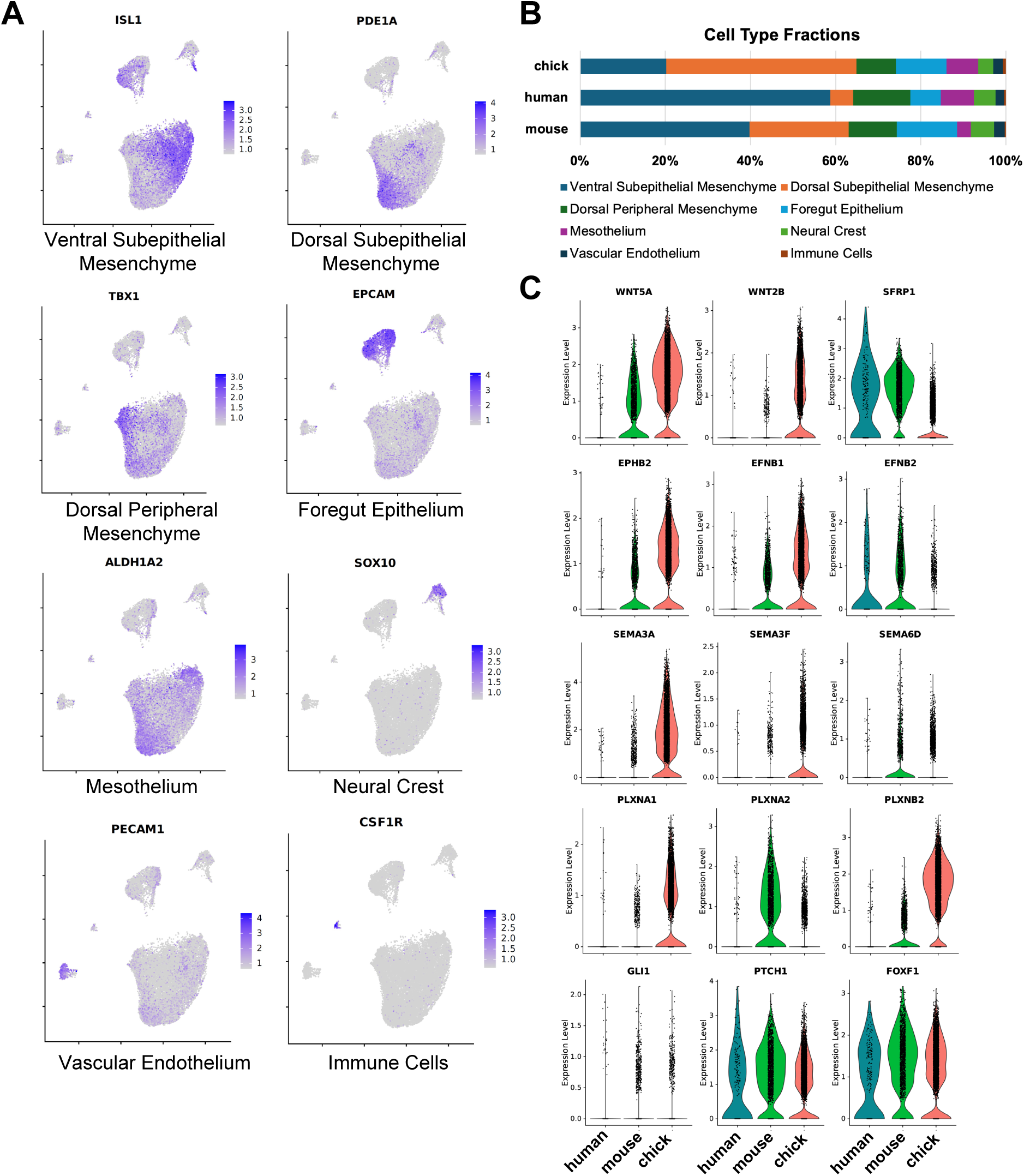
Cell-type annotation and cross-species comparison of integrated human, mouse, and chick foregut scRNA-seq datasets. **(A)** UMAP feature plots showing the expression of representative markers used to annotate the foregut cell populations in Fig. 4A. **(B)** Relative fractions of each cell population among cells from each species in the integrated dataset. **(C)** Violin plots of the expression of selected Wnt signaling (WNT5A, WNT2B, SFRP1), Ephrin signaling (EPHB2, EFNB1, EFNB2), Semaphorin signaling (SEMA3A, SEMA3F, SEMA6D, PLXNA1, PLXNA2, PLXNB2), and SHH response (GLI1, PTCH1, FOXF1) genes in the Dorsal Subepithelial Mesenchyme across human, mouse, and chick foreguts.

## Description of supplementary files

**Video S1**. Two-photon live imaging of an E10.5 nTnG mouse foregut slice, related to Fig. 2E. Timestamp: HH:MM. Scale bar: 50 µm.

**Video S2**. Two-photon live imaging of an E3.75 GFP chick foregut slice, related to Fig. 2E. Timestamp: HH:MM. Scale bar: 50 µm.

**Video S3**. Two-photon activation and live imaging of azidoblebbistatin (left) in the epithelium of an E3.75 chick foregut slice labeled with the membrane dye FLIPPER-TR (right), related to Fig. 3A. Timestamp: MM:SS. Scale bar: 50 µm.

**Video S4**. Long-term two-photon activation of azidoblebbistatin and live imaging of the epithelium of an E3.75 chick foregut slice labeled with the membrane dye FLIPPER-TR, related to Fig. 3B. Timestamp: HH:MM. Scale bar: 50 µm.

